# Decoding the Language of Microbiomes: Leveraging Patterns in 16S Public Data using Word-Embedding Techniques and Applications in Inflammatory Bowel Disease

**DOI:** 10.1101/748152

**Authors:** Christine A. Tataru, Maude M. David

## Abstract

Microbiomes are complex ecological systems that play crucial roles in understanding natural phenomena from human disease to climate change. Especially in human gut microbiome studies, where collecting clinical samples can be arduous, the number of taxa considered in any one study often exceeds the number of samples ten to one hundred-fold. This discrepancy decreases the power of studies to identify meaningful differences between samples, increases the likelihood of false positive results, and subsequently limits reproducibility. Despite the vast collections of microbiome data already available, biome-specific patterns of microbial structure are not currently leveraged to inform studies. Instead, most microbiome survey studies focus on differential abundance testing per taxa in pursuit of specific biomarkers for a given phenotype. This methodology assumes differences in individual species, genera, or families can be used to distinguish between microbial communities and ignores community-level response. In this paper, we propose to leverage public microbiome databases to shift the analysis paradigm from a focus on taxonomic counts to a focus on comprehensive properties that more completely characterize microbial community members’ function and environmental relationships. We learn these properties by applying an embedding algorithm to quantify taxa co-occurrence patterns in over 18,000 samples from the American Gut Project (AGP) microbiome crowdsourcing effort. The resulting set of embeddings transforms human gut microbiome data from thousands of taxa counts to a latent variable landscape of only one hundred “properties”, or contextual relationships. We then compare the predictive power of models trained using properties, normalized taxonomic count data, and another commonly used dimensionality reduction method, Principal Component Analysis in categorizing samples from individuals with inflammatory bowel disease (IBD) and healthy controls. We show that predictive models trained using property data are the most accurate, robust, and generalizable, and that property-based models can be trained on one dataset and deployed on another with positive results. Furthermore, we find that these properties can be interpreted in the context of current knowledge; properties correlate significantly with known metabolic pathways, and distances between taxa in “property space” roughly correlate with their phylogenetic distances. Using these properties, we are able to extract known and new bacterial metabolic pathways associated with inflammatory bowel disease across two completely independent studies.

More broadly, this paper explores a reframing of the microbiome analysis mindset, from taxonomic counts to comprehensive community-level properties. By providing a set of pre-trained embeddings, we allow any V4 16S amplicon study to leverage and apply the publicly informed properties presented to increase the statistical power, reproducibility, and generalizability of analysis.

## 1 Introduction

### 1.1 Microbial survey studies

Microorganisms are biochemically potent entities that influence the biochemistry of surrounding organisms at all ecological scales. Recent findings suggest that resident microbiomes of the human anatomy influence our bodies and minds in ways we have only just begun to understand. Microbiomes have been implicated in the development of diseases of nearly all types, both acute and chronic, infectious and systemic. The vaginal microbiome has been implicated in preterm birth (1), the skin microbiome in acne (2) and eczema (3), and the gut microbiome in a spectrum of diseases including inflammatory bowel disease (IBD) (4–6,6–9), anxiety (10–12), major depressive disorder (13–15), autism (16–20), and Parkinson’s Disease (21–23).

To analyze microbiome compositions, current technology sequences various hypervariable regions of the 16S rRNA gene, which acts as an accessible taxonomic tag to measure the abundances of taxa in a community. Studies using this 16S survey technique have reported incredibly diverse collections of microbes in several systems. Multiple individuals studies, along with the American Gut Project (AGP) (24) and the Human Microbiome Project (25), have invested colossal effort to document that diversity by creating publicly available reference repositories. Amongst these are repositories of stool-associated microbiota that have furthered our understanding of the role of the microbiome in several diseases, especially inflammatory bowel disease (IBD) (4).

Though these and other studies have presented highly relevant findings, 16S microbiome survey studies in general tend to suffer from lack of reproducibility (26,27). Difficulties in reproducibility can be attributed to several technological and analysis-based issues (26,28,29), including three major problems addressed here. First, due to logistical restrictions, especially in human gut microbiome studies where collecting clinical samples can be arduous, the number of taxa considered in any one study often exceeds the number of samples ten to one hundred-fold. Even the largest microbiome studies only include roughly as many samples as taxa analyzed (24,25). As the number of samples necessary to present a statistically sound and reproducible result increases with the number of variables being considered, individual studies with low sample-to-variable ratios risk being underpowered and reporting false positives, especially when effect sizes are estimated to be small (27,30,31).

Second, the most commonly employed analysis techniques assume independence of bacterial species (32–34). In biological contexts, the presence and function of each microbe is deeply dependent on the characteristics of its surrounding neighbors. Differences in microbial function also occur as genes are turned on or off as appropriate for that microbe’s environment at any given time. For instance, Belenguer et al. show that Roseburia strain A2-183 is unable to use lactate as a carbon source except in the presence of Bacteroides adolecentis (35). Because of functional dependence, findings of differential abundance or function of a single species must be considered within its wider context of associated species and environmental factors (36). More specifically, predictive models that differentiate between disease and healthy guts based on microbiome composition in one dataset can rarely be successfully applied to samples from the same patient population collected independently (27).

Third, despite the vast amount of publicly available 16S microbiome survey data, current studies design their data collection and perform their analysis independently, without leveraging the information available via massive sequencing projects such as the Human Microbiome Project (25) or American Gut Project (24).

Navigating the highly related and very large microbiome space can be done by using the information encompassed in publicly available datasets to inform novel dimensionality reduction methods. The goal of this project is to create an unbiased method to project taxonomic data into a lower dimensional space that represents taxa properties. Properties are based on taxa relationships with each other and their environment, are learned from public datasets, and are re-usable for past and future studies. In this context, a property is a pattern that underlies co-occurrences between taxa. The lower dimensional space is learned from public datasets using an embedding algorithm, and allows the integration of patterns from massive datasets into specialized studies to increase reproducibility and statistical power.

### 1.2 Current Methods for Dimensionality Reduction

Currently, most microbiome survey studies focus on differential abundance testing per taxa in pursuit of specific biomarkers for a given phenotype. Often, dimensionality reduction may be performed to reduce the data to a manageable size. For example, taxa may be filtered to consider only the common or very rare, however this approach may filter potentially valuable data. In another approach, taxa can be categorized, or binned, by their phylogenetic relationships (e.g. all taxa that share a family are analyzed as one unit) (37,38). Such binning methods may obscure meaningful biological signal, and are also heavily database dependent not all microbes are clearly classified by taxonomy. Alternatively, taxa can be clustered based on the similarity of their 16S rRNA gene, which has been used as a proxy for evolutionary relatedness (39). However, in this approach, clustering may hinder comparisons across studies, and may result in biologically unfounded taxonomic units (28). Such taxonomic count-based methodologies, while they have led to interesting and crucial discoveries in stool-associated microbiome surveys, assume that differences in individual species, genera, or families can be used to distinguish between microbial communities and ignore community-level action between and among species.

Rather than searching for individual biomarkers, ordination may instead be used to reduce data dimensionality and identify broad patterns in microbiome compositions between samples. Samples, each represented by a vector of taxa, can be projected into a lower dimensional space using a wide array of ordination techniques including principal component analysis (PCA) (40) and multidimensional scaling (41). Broadly used, ordination has played a critical role in associating microbial structure with specific features or phenotypes of interest, but has also proven to be overly sensitive to normalization and study bias (e.g. technological noise, DNA preparation protocol, sequencing error) (42).

Several studies have attempted to integrate phylogenetic or edit distances between 16S gene variants. Woloszynek et. al represent each 16S sequence by the set of k-length nucleotide sequences (k-mers) it includes, and embed those k-mers to create a vector representation of each sequence (43).

Finally, we may use taxa counts to try to estimate parameters for an underlying distribution from which taxa are drawn. Sankaran et. al model taxa as units drawn from a latent Dirichlet multinomial mixture distribution 36). This method aptly describes samples by assigning topic distributions to them, but does not directly relate taxa to each other, and does leverage available public data.

### 1.3 Current Study Proposal

While compelling, the dimensionality reduction methods described above do not consider taxonomic relationships within a biological context, or make use of information already available from previous datasets. By integrating previous studies and subsequently putting 16S rRNA gene into context, our study proposes to describe inherent properties of a microbial communities in a manner consistent with their functional utility in their environmental context.

To deduce the above-mentioned properties, we turn to embedding techniques from natural language processing. The use of natural language methods in microbiome analysis is not new. As noted by Sankaran et. al (36), there exist some easily drawn parallels between natural language data and microbiome data, namely that documents are equivalent to biological samples, words to taxa, and topics to microbial neighborhoods. Just as a book may be defined by the aggregate topics it discusses, a microbial environment may be defined the neighborhoods or communities it contains.

There is another connection between words and microbes not currently discussed in the literature, and that is the capacity of both entities to be described by a finite set of discrete, characteristic properties. For instance, the word ‘apple’ in English can be defined as an edible, red, non-gendered, crunchy, object. Similarly, the species Clostridium difficile can be defined as a spore-forming, infectious, spindle-shaped bacteria. While it would be difficult to distinguish between a recipe book and a magazine of food reviews by enumerating differences in the occurrence of individual words, differentiating the two becomes simple if we select appropriate properties. While both media use words that have high scores in the property “edibility”, the recipe book also uses words that have a high declarative score, like ‘cut’, ‘wash’, and ‘prepare’, while the food review uses words that have high descriptive scores, like ‘fantastic’, ‘delectable’, or ‘abysmal’. Just as the properties of “declarative” and “descriptive” allow us to differentiate texts more effectively, property-based analysis of microbiomes allow us to distinguish between two microbial scenarios more easily than individual taxa counts. Analysis on the level of properties thus provides a more accurate and generalizable representation of the data’s structure.

In this study, the properties mentioned above were learned from patterns in a large microbial dataset provided by the American Gut Project (AGP). An unsupervised embedding algorithm developed for natural language processing called GloVe (44) was applied to over 15,000 AGP samples to learn an embedding space by quantifying co-occurrence patterns between taxa. The resulting set of embeddings transforms human gut microbiome data from thousands of taxa counts to a property space of only one hundred to seven hundred variables. We quantify the quality of the properties by predicting the Inflammatory Bowel Disease (IBD) status of samples using properties, normalized taxonomic count data, and principal component analysis. We show that predictive random forest models trained using property data are the most accurate, robust, and generalizable, and that property-based models can be trained on one dataset and deployed on an independent one with positive results. Strong correlation between learned properties and annotated metabolic pathways allow us to implicate both known and new metabolic pathways in IBD such as steroid degradation, lipopolysaccharide biosynthesis, and various types of glycan biosynthesis. Lastly, by projecting taxonomic data into property space, the scientific community can integrate patterns from massive public datasets into specific, targeted studies. Analysis in property space means models requires fewer samples to produce robust results, and exploratory studies simultaneously gain increased power and decreased risk of spurious associations.

We not only advocate the use of this method, but also propose to shift the analysis paradigm from a focus on taxonomic counts to a focus on comprehensive properties that more completely characterize microbial community members’ function and environmental relationships. The human gut microbiome has the potential to be used as a low-cost environmental barometer for the diagnosis and monitoring of disease, but first we must prioritize model reproducibility and move beyond the concept of the taxonomic unit.

## 2 Results

### 2.1 Model performance

In order to determine the value of the set of embedding produced by GloVe, we tested the performance of classifiers built using GloVe embedded, PCA transformed, and non-embedded normalized count data. We evaluated two main performance metrics in predicting the IBD status of the host: area under the receiver operating curve (AUROC) and area under the precision-recall curve (AUPR). The receiver operating curve plots true positive calls against false positive calls. The higher the AUROC, the more confident you can be that a positive prediction by the classifier is correct. The precision-recall curve plots the precision, how confident you are that a positive call is correct, against recall, what percentage of the positive samples in the dataset were identified. A high AUPR means the classifier is able to identify most of the positive samples without making too many false positive calls. Both curves plot these values over a range of decision thresholds. For both metrics, a value of 1 is a perfect classifier.

### 2.2 Pick optimal number of properties to define a community

We found random forest classifiers trained using GloVe embedded data produce a significantly higher average area under the Receiver Operating Curve (AU-ROC) across all choices of hyperparameters and number of dimensions (Fig 2) than non-embedded data and PCA-embedded data (p *<<* 0.05, rank sum test). Notably, embedded data consistently produces better results with far fewer features than taxonomic counts. The use of fewer features makes the model less likely to overfit the data and more likely to be reproducible. We run all future tests using 100 properties, as models trained with 100 properties show the most consistently high performance and small variance across all hyperparameter choices.

**Figure 1:**
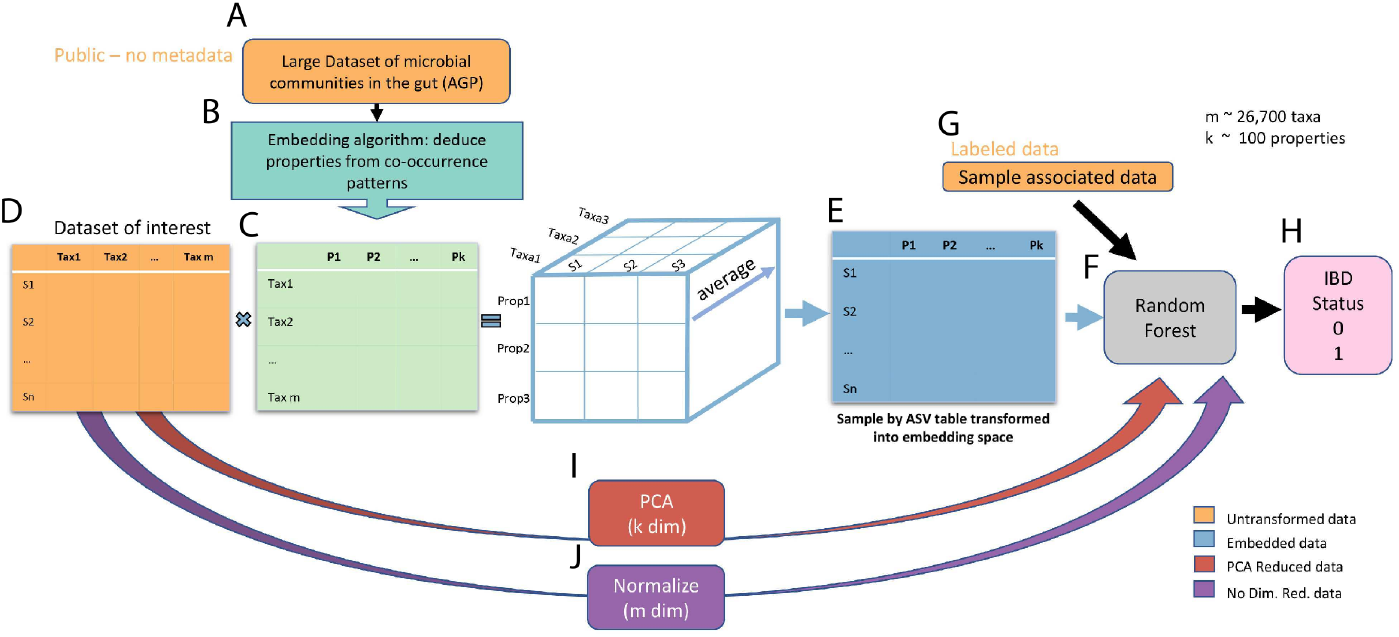
Workflow of data transformation to prediction of host phenotype. First, taxa-taxa co-occurrence (binary) data from the American Gut Project (A) are input into the GloVe embedding algorithm (B) to produce a taxa (Amplicon Sequence Variant or ASV) by property transformation matrix (C). Then, we take the dot product between a sample by taxa table of interest (D) and the transformation matrix (C) to project that table into embedding space (E). This table is used to train a random forest model (F) along with sample associated lifestyle and dietary information (G) to predict the IBD status of the host (H). As points of comparison, random forest models are also built without embedding, after transforming the same sample by taxa table (D) using PCA (I) and normalizing (J).

**Figure 2:**
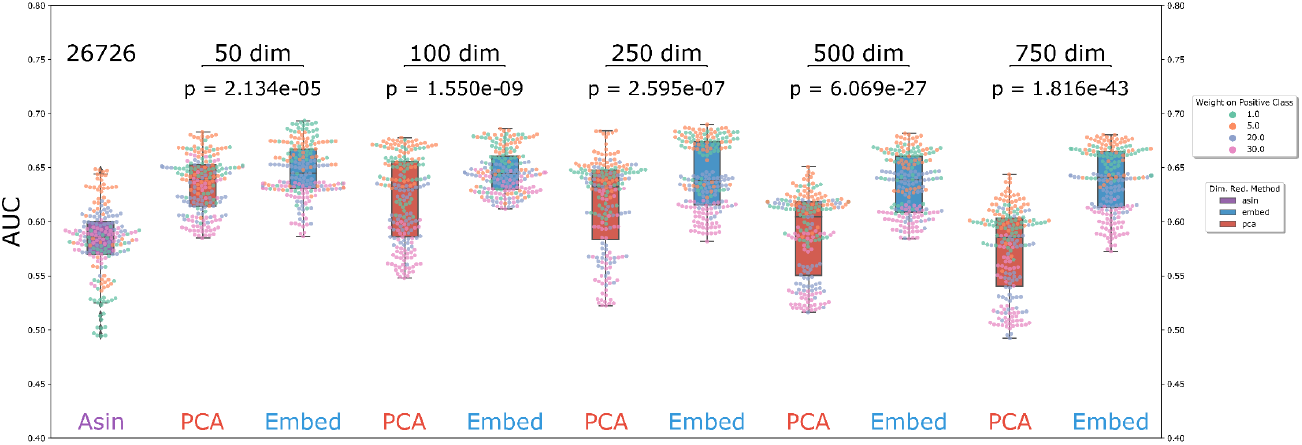
Transforming ASV tables into GloVe embedding space before training a model produces more accurate host phenotype predictions (IBD vs. healthy control) and makes models more robust to hyperparameter choice. Each point represents a triplet of choices for number of trees, depth of each tree, and weight on a positive prediction of IBD in a random forest model. Each model was trained on the data input type indicated by color (Normalized, non-embedded counts is purple, PCA embedded data is pink, and GloVe embedded data is blue). Models trained on GloVe embedded data produce higher ROC AUCs with less variance across hyperparameter choice.

### 2.3 Models built with embedded data perform better on a held out test set

We then train three separate models on the training portion of the AGP dataset, and test each model on a held out portion of the same dataset that has been used neither for model nor embedding training (Fig 3 panel A). Each model uses a different data input type, GloVe embedded, PCA-transformed, or non-embedded normalized taxa counts, and has hyperparameters optimized using cross-validation over the training set. We see comparable performance between the classifier using GloVe embedded data and the other two methods (Fig 3 panel B). The model with non-embedded data, which uses 26,739 features, has an area under the Receiver Operating Curve (AUROC) of 0.79 and an area under the Precision-Recall curve (AUPR) of 0.46 (Fig 3, panel B.1). In contrast, the model using GloVe embedded data, which uses only 113 features, has a higher AUROC of 0.81 but slightly lower AUPR of 0.44 (Fig 3 panel B.2). A 200-fold decrease in number of features used results in little change in relevant performance metrics. In comparison, the model using PCA-transformed data with 113 features performs only slightly worse, with an AUROC of 0.77 and an AUPR of 0.42 (Fig 3 panel B.3)

**Figure 3:**
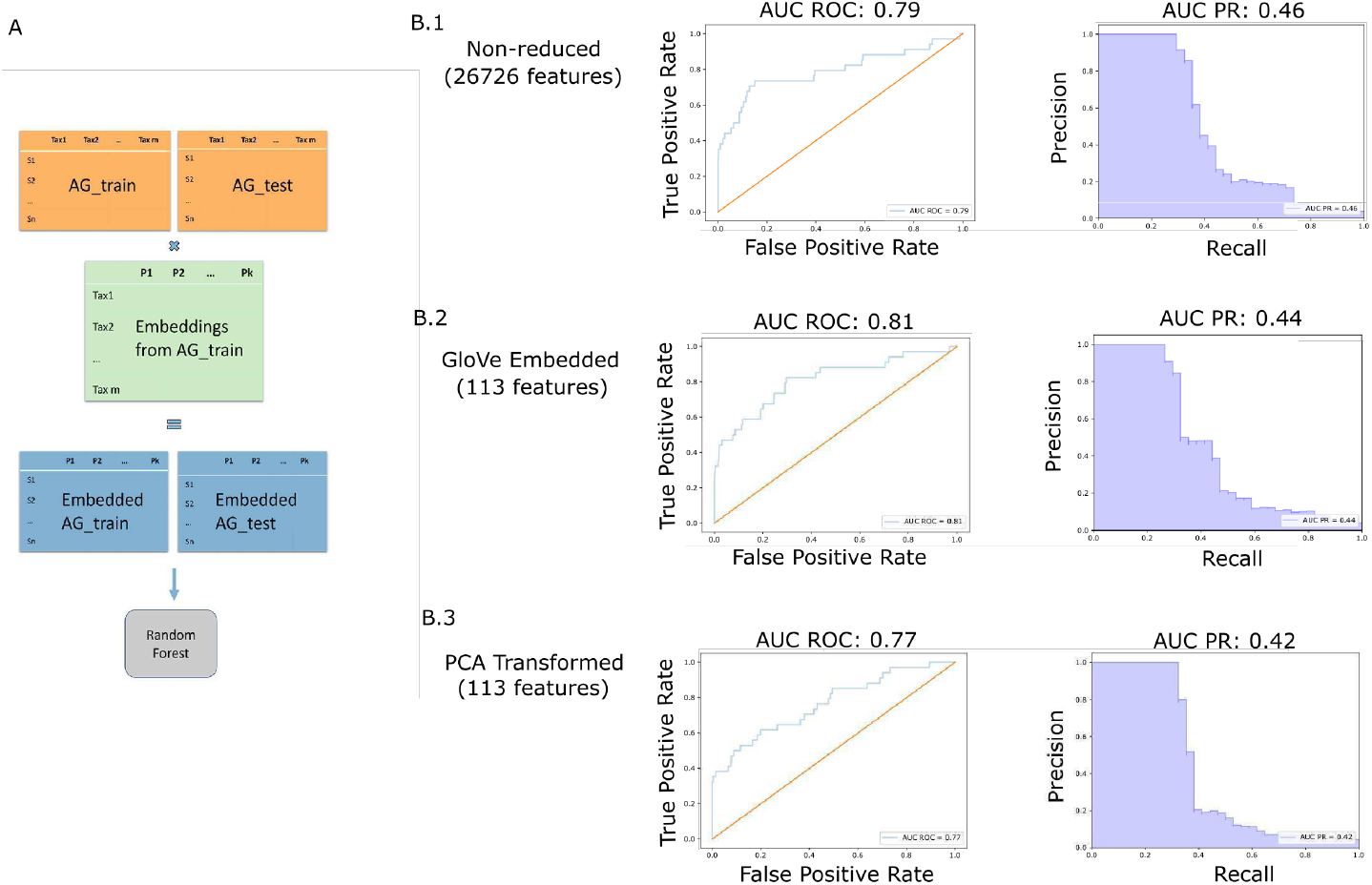
Embeddings trained on American Gut training set, model trained on American Gut training set, model tested on American Gut held out test set (A). Models trained on GloVe embedded data have higher ROC AUC but slightly lower Precision-Recall AUC on a held out test set (B)

### 2.4 Properties are generalizable to independent stool-associated datasets

We find that GloVe embedded data generalizes to a completely independent datasets, and significantly improves performance when fewer than 400 training samples are available. Using data from Halfvarson et. al (8), we train random forest classifiers on gut microbiome data to differentiate between IBD vs. healthy control (Fig. 4A). Again, we train classifiers using normalized count data, PCA-embedded data, and GloVe embedded data, and optimize over hyperparameters using cross-validation for each model independently. To test the effect of training set size on performance outcomes, we train models using from 50 to 450 samples in the training set, and the rest in the test set. In this dataset, we have 564 samples from 118 patients and 17, 775 Amplicon Sequence Variances (ASVs). We do not include any associated metadata; predictions are made solely based off of the microbiome compositions.

**Figure 4:**
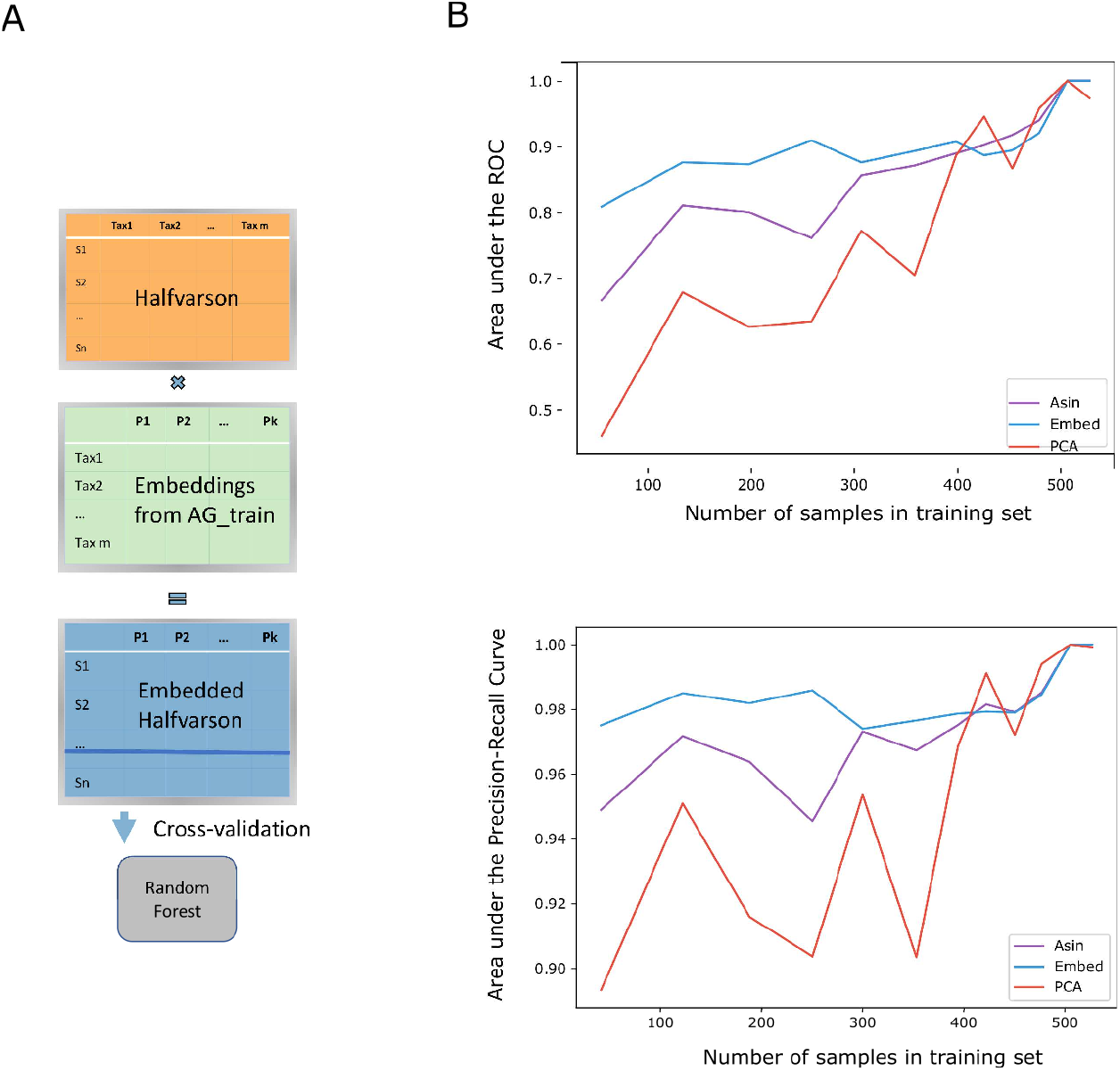
Embeddings trained on American Gut data, model trained and tested on Halfvarson dataset (A). Transforming microbiome data into GloVe embedding space prior to model training produces more accurate models despite smaller training sample sizes (B).

It is important to note that the transformation matrix that puts the query dataset into embedding space is trained exclusively on American Gut Project data, and is therefore completely independent of the query dataset. Despite the fact that properties were learned using the American Gut data dataset exclusively, we see better embedding model performance on the independent set from Halfvarson et. al (8) (Fig. 4B). In particular, we see that as the number of training samples becomes smaller, embedding-based models are able to maintain high AUROC (Fig. 4B.1) and AUPR (Fig. 4B.2) while models based on PCA-transformed data (100 features) and non-dimensionality reduced models (17,775 features) cannot. When large numbers of training samples are available, all methods perform comparably, but only embedding-based models perform well at middling to low (*<* 400) sample sizes.

The patterns learned by the GloVe algorithm from the American Gut data generalize to improve classification performance on an independent dataset. Theoretically, classification accuracy of any host phenotype relating to the gut microbiome could be bolstered by first embedding the input data before model training.

### 2.5 Models that use properties are generalizable to independent datasets

In the above experiments, all models were trained on the same datasets they were tested on, using cross-validation and a held-out test set. Now, we trained a model on the American Gut data and tested it on the Halfvarson data (Fig 5A). More so than a hold-out test set, this allows us to test the feasibility of deploying a model for diagnosis and monitoring of IBD. Two models were trained, one using normalized taxa counts and the other taxa counts embedded in property space. In this case, only microbiome data and no sample-associated data was included. Hyperparameters that gave the highest F1 score on American Gut data were selected, and the trained model was directly applied to the independent dataset without re-tuning hyperparameters or decision thresholds. Both models trained on American Gut taxa count and American Gut embedded data had a precision of 1, meaning that a positive IBD prediction was correct 100 percent of the time. However, the model trained on taxa counts had a recall of 0.02, meaning that only 2 percent of the samples from patients with IBD were positively identified. In contrast, the model trained on embedded data recovered 26 percent of samples from patients with IBD. While the model trained on taxa counts was in no way generalizable to another dataset, the model trained on data in property space was able to make accurate predictions on a completely independent dataset (Fig 5B). This finding demonstrates that in this case, models built from embedded data can generalize to outside data while models built from taxa abundance information cannot.

**Figure 5:**
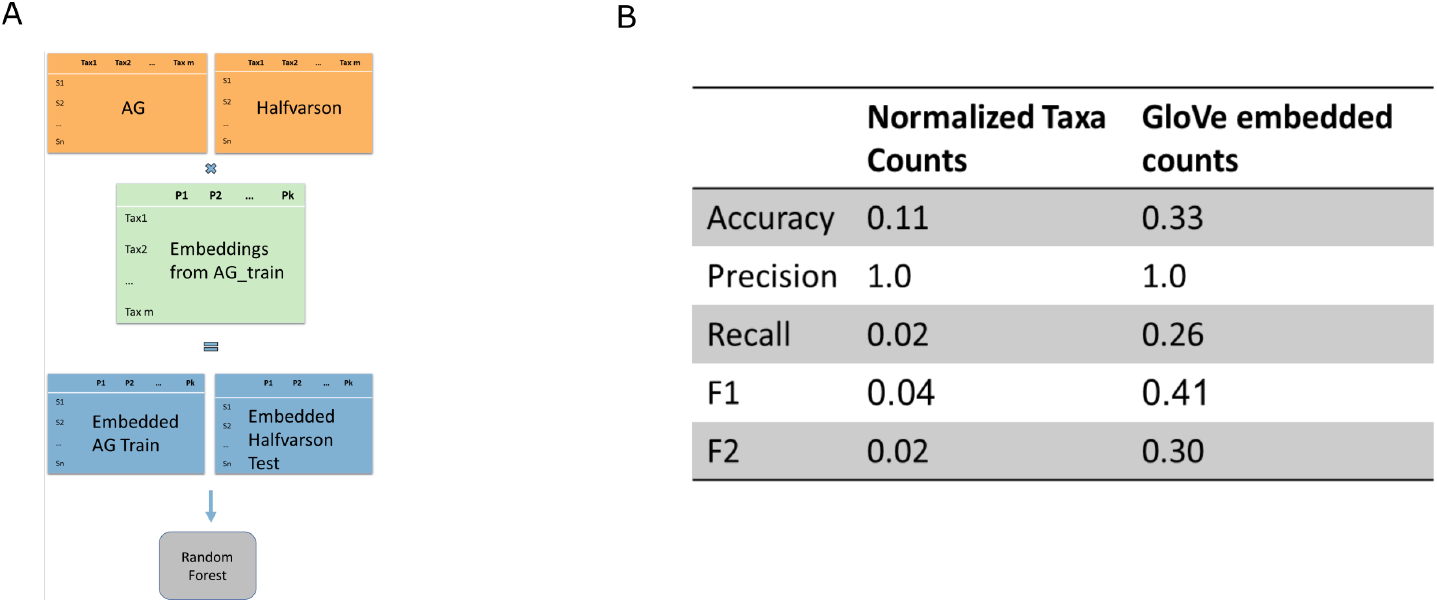
Models and embeddings trained on American Gut data and tested on Halfvarson data (A). Model trained on properties far outperforms models trained on taxa counts (B).

### 2.6 Distances in embedding space roughly correlate with phylogenetic distance

Taxa close together in embedding space have similar co-occurrence patterns. We expect that phylogenetically close taxa are more likely to fill the same ecological niches than are unrelated taxa. We therefore expect a slight but not extreme correlation between phylogenetic distance and distance in embedding space. Using a Mantel test (45), we do observe a low (coef = 0.12) but significant (p = 0.001) correlation between the two distance metrics, with more granularity available when comparing taxa in embedding space. This finding demonstrates that co-occurrence patterns as captured by embeddings are a more sensitive distance metric than phylogeny (Fig 6).

**Figure 6:**
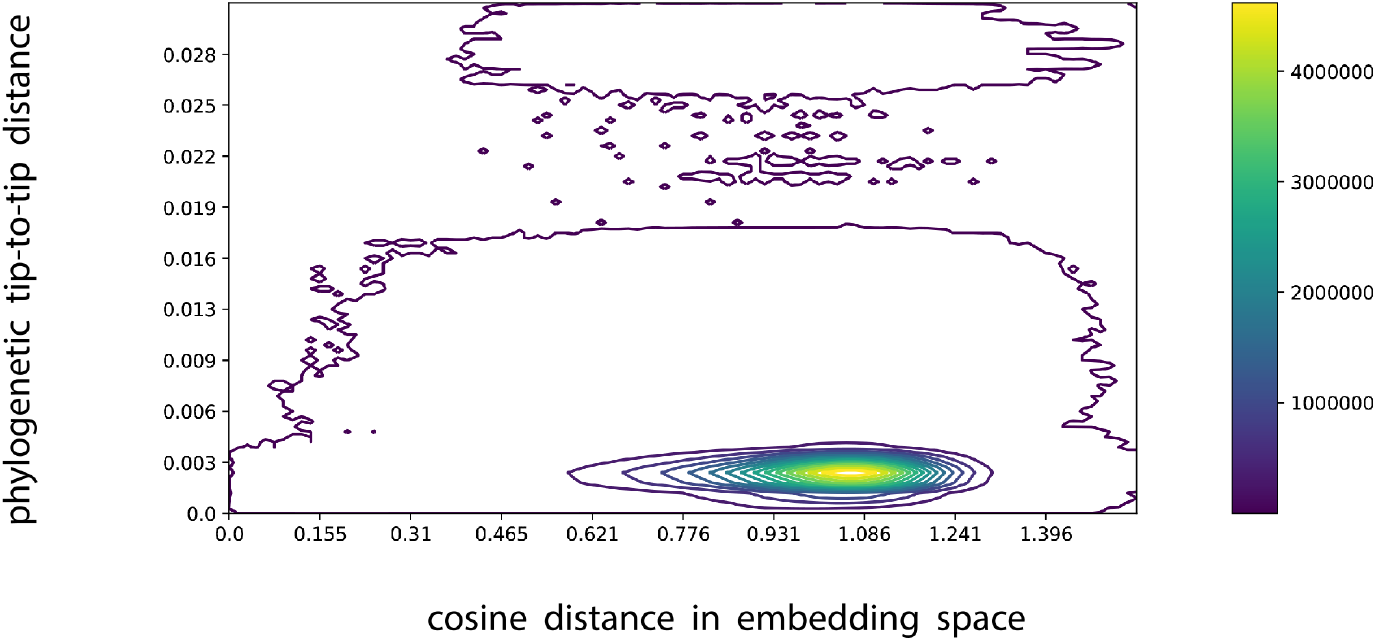
The contour plot shows that distances between pairs of taxa in GloVe embedding space roughly correlate with distances between those taxa in phylogenetic space (A). A lighter color signifies a higher density of taxa pairs. There is more granularity along the embedding space axis, implying that related taxa are more easily distinguished from each other in embedding space than they are phylogenetically. A Mantel test shows a low slope but very statistically significant correlation between the two distance metrics (p = 0.001)

### 2.7 Relationship with Metabolic Capacity

We chose to preserve taxa co-occurrence patterns in embedding space because we hypothesize that those patterns are driven by taxa functionality in an environment. As such, we evaluate the possibility of a connection between annotated genetic capacity to express metabolic pathways and the properties that make up embedding space. First, we find each Amplicon Sequence Variant’s (ASV) nearest neighbor in the KEGG database (46) using Piphillian (47), and use the KEGGREST API (48) to determine which pathways are present in that ASV’s genome. This results in an ASV by pathway table where there are 11,893 ASVs with near neighbors in the database, and 148 possible metabolic pathways. Then, we identify the significantly correlated metabolic pathways for each property in embedding space. A permutation test is used to simulate a null distribution of maximum correlations per embedding property and determine significance. We find that every property significantly correlates with at least 1 annotated metabolic pathway. Suppl. Table 1 shows each dimension and its significantly correlated metabolic pathways; each dimension has significant correlation with 3 to 57 pathways. We see that the magnitude of correlations between embedding dimensions and metabolic pathways are far greater in the GloVe embedding case than in the PCA-transformed case (Fig 7). Additionally, none of the correlations between PCA dimensions and metabolic pathways are significant under a permutation test after multiple hypothesis correction (Suppl. Fig. 1). This suggests that the properties learned by the GloVe algorithm based on co-occurrence patterns between taxa may actually reflect the metabolic capacity of those taxa.

**Figure 7:**
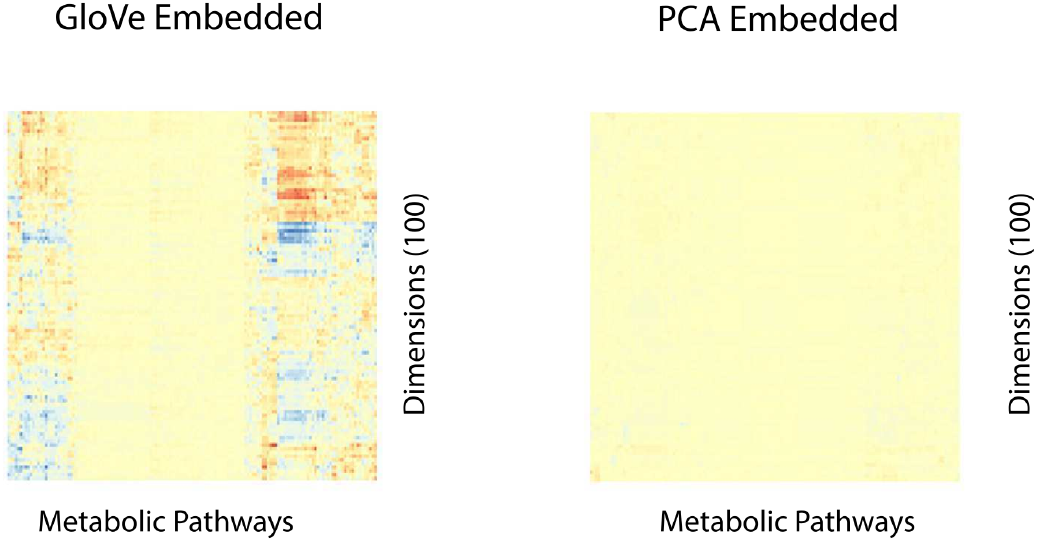
Dimensions in GloVe embedding space correlate with some metabolic pathway annotations (A), but dimensions in PCA embedding space do not (B). Each column in each heat map represents a metabolic pathway from KEGG (e.g. ko00983). Each row is a dimension in either GloVe or PCA embedding space.

### 2.8 Interpreting the predictive model for IBD with metabolic pathways

In order to explore the implications of properties and metabolic pathways for IBD, we calculated an association score (see method section 4.8) between each property and a positive IBD prediction. The full tables of the most predictive dimensions, their associated metabolic pathways, and their direction of influence on the prediction can be found in Suppl. Table 2 (AG) and Suppl. Table 3 (Halfvarson). First, we identified those properties strongly associated with a positive IBD prediction (association score above 8 in both Halfvarson and AGP datasets). We then selected all metabolic pathways significantly correlated with more than one of those highly relevant properties. In this way, we identified 45 metabolic pathways of interest for IBD (Suppl. Table 4). The pathways fall broadly into 9 main categories according to the KEGG Brite database: steroid metabolism, lipid metabolism, glycan biosynthesis, amino acid metabolism, antibiotic synthesis and resistance, bacterial pathogenic markers, metabolism of terpernoids and polyketides, cell motility and cellular community formation, and xenobiotics biodegradation/other metabolic function.

### 2.9 Explaining the variance in properties

Lastly, we sought to determine how much of the information contained in properties can be recapitulated by looking at the above described annotated metabolic pathways, and how much was unique to each property. For each property, we use a linear regression to predict the property values per taxa from the pathway presence/absence per taxa. We report the *r*^2^ statistic per property, and find that metabolic pathways can explain a maximum of 36 percent of the variance in one property, and a minimum of 11 percent of another. This means that, while there is a strong correlation between properties and annotated metabolic pathways, most of the information contained in properties are not represented by annotated information (Suppl. Fig. 2)

## 3 Discussion

In a data-driven field dominated by small sample sizes and large variable spaces, it is necessary to employ some form of dimensionality reduction. Currently, this is done by filtering by taxa prevalence, clustering based on phylogenetic proximity, or is not done at all. We present here a method to leverage massive public datasets to learn an embedding space that represents the latent properties driving taxonomic abundances. By shifting the paradigm of analysis from taxonomic counts to community-level microbiome properties, we enable more holistic, comprehensive analysis that accounts for taxonomic relationships while simultaneously simplifying the data.

We demonstrate that we can learn a fecal microbiome property space that is more apt at predicting the IBD status of the host than non-reduced and pca-reduced spaces, and remains accurate even at low training sample sizes. We also present a classification model trained on property data that generalizes well between datasets, where models based on taxonomic counts do not. We lastly define the relationships between embedding space and known metrics used to explore microbiomes like phylogenetic distance and metabolic pathway genetic capacity.

### 3.1 Properties

Embedding is a technique used ubiquitously in machine learning, especially in natural language processing (49–51). Embedding algorithms take discrete units of data (e.g. words or taxa) and embed them into a vector space, preserving proximity between the units based on any metric that can compare two units. In the case of embedding taxa, possible metrics include phylogenetic distance, genome similarity, or morphology: in this paper the chosen metric to determine proximity between units is patterns of co-occurrence. The embedding algorithm used in this paper is GloVe, an algorithm designed for word processing (44). Using this algorithm, two taxa that occur with similar sets of other taxa at similar frequencies should be close in embedding space, and two taxa that are found in the presence of different neighbor sets should be far from each other. To visualize this, we return to the analogy of word analysis. Two words, “apple” and “banana”, are close to each other in embedding space because they tend to occur with similar sets of words like “eat”, “fruit”, “tasty”, and “smoothie”. Likewise, the words “king” and “marshmallow” tend to occur in different contexts; “king” is most often found in the company of words like “politics”, “throne”, and “empire” while “marshmallow” is found with words like “toddler”, “fluffy”, and “scrumptious”. Note that there are two ways words may be close in embedding space. First, words may directly co-occur frequently, like the words “apple” and “banana”. Instead, words may be synonyms, which do not often co-occur directly with each other, but instead co-occur with similar patterns, like “large” and “huge” both being used to describe giants, mountains, and appetites. Returning to the concept of embedding taxa, we may use embeddings to discover relationships both between taxa that work together directly, and between taxa that are synonymous and likely fill the same niche.

Once proximity in embedding space has been established, the data can immediately be used to improve modeling efforts. Subsequently, conceptual properties can be assigned to the learned dimensions by observing which entities have similar values in any given dimension. If “strawberry”, “cookies”, “cake”, and “ice-cream” all have high values in one dimension, and “mud”, “medicine”, and “brussel sprouts” all have low values in that same dimension, we may call that dimension the “delicious” property.

We have shown that embedding an ASV table into property space using GloVe integrates patterns from public data into modeling efforts, producing more accurate diagnostics while decreasing data dimensionality. Classifiers built after transforming data in this way are more robust, and the same embeddings generalize to improve the accuracy of classifiers built from completely independent datasets. Properties also allow models trained on one dataset to be applied to another independent dataset with positive results.

In addition to improving classification accuracy for IBD, the embeddings quantify and simplify the microbial landscape of gut microbiomes. Rather than considering a microbiome as a collection of bacterial counts, all of which are mostly independent, we propose to describe a microbiome as a vector of values for the relevant properties. Consider the example of distinguishing the recipe book from the food magazine; reducing each into property space allows us to clearly see the differences in declarative and descriptive word usage rather than counting the number of times the words “spinach”, “tomato”, and “bowl” were each used. Because these properties are learned from the data directly, they are much less biased than manually engineered features. Analysis performed on this latent property space is likely to be much more robust to variations in datasets, addressing the problem of irreproducibility currently plaguing 16S microbiome studies (26,27).

### 3.2 Biologically driven dimensional reduction

We use unsupervised learning to define an embedding space where taxa proximity represents similarity in co-occurrence patterns. Unsupervised learning limits the human decision-making bias in property definition, but also produces unlabeled properties whose interpretation is not immediately obvious. We hypothesize that co-occurrence patterns are driven by taxa function like metabolism, synthesis of secondary metabolites, and secretion of antimicrobial products. We show in our pathway analysis that property distributions in fact do correlate significantly with metabolic pathways. Therefore, the learned property space is likely informed by taxa function from within a biological context. Some elements of property space may also be informed by other factors, such as geography or diet commonalities between groups of people, and this should be explored further.

### 3.3 Annotation Independent

While we have explored the associations between embedding properties and the annotated quantities of genetic potential, the power of this embedding technique is that it does not rely on annotations of known taxonomic groupings or full genomes in order to improve prediction accuracy of host phenotype. Because any ASV that has been observed during embedding training can be embedded, it is possible to describe the properties of uncultured and unannotated ASVs, and include this information in a classifier. The transformation into embedding space requires only an ASV table, and uses no sample associated data like lifestyle variables or diagnoses.

### 3.4 Implications for IBD

We were able to identify 9 main categories of KEGG BRITE pathways that were significantly correlated with properties associated with IBD (Suppl. Table 4). Among these pathways, both steroid metabolism and biosynthesis were found to be associated with IBD. Steroids are a well-known and commonly utilized treatment for patients with active Crohn’s disease (52). Enrichment in steroid metabolism in the gut microbiome could be reflective of an increase in steroid availability due to treatment.

Several pathways belonging to the rather broad BRITE category of “other metabolic function” have already been well explored and characterized in the literature as related to IBD. Toluene degradation (KEGG pathway 00623) was found to be increased in both Crohn’s disease (CD) and Ulcerative Colitis (UC) samples in a microbiome survey meta-analysis (53). Components of the benzoate metabolic pathway, including fluorobenzoate degradation (KEGG pathway 00364), were associated with IBD severity in a treatment-naive cohort with CD (54). Analysis of inflamed gut lining mucosa in patients with IBD also found decreased ascorbic acid content (KEGG pathway 00053)(55) All of these pathways, along with dioxin degradation, inositol phosphate metabolism, and lipoic acid metabolism, were associated with an IBD prediction in our model.

We also found multiple glycan biosynthesis pathways correlated with predictive IBD properties (KEGG pathways 00511, 00514, 00515, 00601). In particular, bacterial glycosphingolipid biosynthesis, a pathway which has antiinflammatory effects when produced by the host epithelial cells (56), was found to be associated with IBD in our model. We speculate that this may indicate a shortage of glycosphingolipids in the gut environment, exacting positive selection pressure on microbes that can produce their own. Lipopolysaccharide biosynthesis and multiple types of O-glycan biosynthesis were also implicated in our model, all of whose association with IBD has been explored, briefly, in the literature (57–59). Given its importance and consistency in our predictive model, this group of pathways may warrant further exploration.

### 3.5 Limitations and future expansion of the work

While embedding Amplicon Sequence Variants (ASVs) affords the benefits to classification and interpretation previously discussed, it relies heavily on the definition of a “biologically meaningful unit” which will then be embedded. For the sake of between-study replicability, we choose to measure the co-occurrence patterns of ASVs (28) as a base unit. It may, however, prove more informative to define a biologically meaningful unit in another way. For example, perhaps ASVs clustered at a 99 percent threshold more accurately capture meaningful patterns in co-occurrence. We may also consider a variable threshold that is more representative of common ancestry on a phylogenetic tree and aggregate based on clade architecture before embedding.

Additionally, the presented set of embeddings was constructed using only the forward reads from the American Gut dataset, as reverse reads were not provided in the EBI database. Future embeddings constructed from full length V4 or other 16S hypervariable regions will likely provide more accuracy and specificity. New embedding transformation matrices would need to be trained for each new biome or segment of 16S gene being explored.

In its current form, the algorithm does not make specific considerations for differences in sequencing depth, which affects how many taxa can be observed in a given sample. Future iterations of this method could include weights such that the observed absence of taxa in a sample with a large number of reads is weighted more heavily than the absence of taxa in a sample with fewer reads.

While the construction of embeddings is not affected by the inconsistency of self-report data, the accuracy of the classifier may be. In this study, we considered only a self-reported medical professional diagnosis to be accurate, and rejected any self-diagnosis reports. While it is possible that classifier performance would change with the inclusion of more liberal diagnostic criteria, the strict diagnosis definition successfully generalized to an independent dataset, which was not self-reported (8).

Properties in embedding space have strong associations with metabolic pathway potentials, but it remains unclear whether they truly represent the expression of those pathways. Future development could also consist of integrating multi-omics datasets available in other studies, including the Human Microbiome Project. Wet lab validation of these hypothesized property-metabolic expression associations would verify the ability of GloVe embeddings to predict metabolic expression from 16S data. This would allow for the integration of metabolic data from all observed taxa, not just those few whose full genomes are available in databases.

It might be possible to use the embedding space to identify taxa that form stable communities together - taxa that are close in embedding space may stabilize each other in culture and in vivo. Through mechanisms like cross-feeding, joint nutrient acquisition, and other cooperative behaviors, microbes may form groups that are more versatile and secure than the individual species on their own. Taxa near each other in embedding space, if they are not directly interacting, may have synonymous functions in their respective communities. By clustering and categorizing microbes by their respective roles, we may gain insight into which bacterial populations secure one another’s stability. Particularly, the relationship between phylogenetic distance and distance in embedding space may be of interest. Microbes that are very closely related to one another through evolution but have very dissimilar co-occurrence patterns may be particularly predictive of their environment, as they have specialized quickly and efficiently. It may be that different variable regions better capture the co-occurrence patterns of taxa, and so are more representative of taxonomic relationship to the environment.

Lastly, the embedding framework can be applied to any system or base unit of interest. It may be particularly illustrative to embed genes from metagenomic datasets instead of taxa. This would allow us to determine mathematical representations of the context of each gene, as well as to glean the robustness and reproducibility benefits from dimensionality reduction for metagenomic data. As always, appropriate benchmarking and exploratory analyses will be necessary to determine the appropriate use cases for this technology.

### 3.6 Conclusion

By integrating patterns from public datasets into individual survey studies, we bring the increased statistical power and generalizability of results of meta-analyses into each independent study. While this work shows the value of an embedding framework for predicting IBD from the gut microbiome, this same framework can be leveraged in any environment with enough data and for any predictive problem of interest. Furthermore, we assert that analyses that define microbiomes by their latent properties instead of by their taxa member list are more informative, reproducible, and relevant to the macroscopic world.

## 4 Material and Methods

Code available at: https://github.com/MaudeDavidLab/embeddings

### 4.1 Embeddings: GloVe algorithm

We used the GloVe algorithm (44) on ASVs to generate embeddings. Briefly, the embedding algorithm (Figure 1B) learns taxa representations that maintain patterns in co-occurrence between pairs of taxa, and was used to learn properties of microbial context. In this algorithm, the metric to be preserved is a function of P_*i*_*k*/*P*_*j*_*k, theprobabilitiesofco occurrenceoftaxaiandjwithk.V ariablesiandjarethetaxabeingrelated, andkisa dimensionalspace, wherexischosenbytheuser.Thex-dimensionalspaceissharedacrossalltaxa, andthuseachdi50, 100, 250, 500, and 750.Embeddingswerelearnedon85percentofthedata, which15percentofsamplessetasidef*

### 4.2 Transformation into embedding space

In 16S survey studies, each sample is represented by a vector of its taxa abundances. Thus, we transform samples into embedding space simply by taking the dot product between each sample’s taxa vector and a taxa’s property vector. This gives an average of property values weighted by taxa abundance. We consider two ASVs the same taxa if they are at least 97 percent similar and align with an e value less than 10^−^29.

### 4.3 PCA transformation

Predictive model performance using embedded data was compared against models trained on data transformed with Principal Coordinate Analysis (PCA). PCA is an ordination technique that projects samples into lower dimensional space while maximizing the variance of the projected data (60).

### 4.4 Random Forest Predictions

The value of the embeddings was evaluated by success at predicting host IBD status using a random forest model (60). The model was built using Python sci-kit-learn, and hyperparameters for the depth of tree, number of trees, and weight on a positive prediction were selected using 10-fold cross validation on the training set. A different model with different hyperparameters was built for each data type, normalized taxonomic abundances, PCA embedded abundances, and GloVe embedded abundances. Counts were normalized by applying an inverse hyperbolic sin function. Models also included self-reported sample metadata such as exercise, sex, daily water consumption, probiotic consumption, and dietary habits. Models were evaluated by their performance, namely area under the receiver operating curve, on the held out test set of 15 percent of samples.

### 4.5 Correlations with KEGG Pathways

For each ASV, we find its closest match, thresholded at 97 percent similarity, in the KEGG database using the software Piphillan (47). Each possible metabolic pathway then gets assigned a 0 if it is absent or a 1 if it is present in that nearest neighbor’s genome.

Limiting the following analysis to include only those taxa that had near neighbors in the database, for each of the properties in embedding space, we find its maximally correlated (absolute value) metabolic pathway. Then, to ascertain whether those correlations were significant, we applied a permutation test (61). We constructed 10,000 null pathway tables by permuting the rows of the original pathway table. We repeated the above procedure, finding the maximally correlated pathway for each of the embedding dimensions in each of the null pathway tables. This results in 10,000 maximum correlation values per embedding dimension, which form a null distribution for each embedding dimension. The significance of the statistic in a permutation test is calculated as the number of times a maximum correlation in a null pathway table was more extreme or equal to the maximum correlation actually observed. Dimensions (columns) in both GloVe transformation matrix and PCA rotation matrix space are normalized to mean 0 and variance 1 to account for differences in scales between the two spaces. We report both the maximally correlated pathway for each property, all of which are significant, and also all significantly correlated pathways per property.

### 4.6 Calculating Phylogenetic Distances

We produced a Multiple Sequence Alignment and subsequently a phylogenetic tree of all ASVs, using Clustal W2 (62) multiple alignment and phylogeny creation software. The tip-to-tip phylogenetic distances were then calculated between every pair of taxa using the dendropy python package (63).

### 4.7 Explaining variance of properties with metabolic pathways

For each property, we set up a linear regression where the property values per taxa are the response variable, and the pathway presence/absence for each taxa are the independent variables. The *r*^2^ statistic is reported to assess the variance in property values explainable by the presence of annotated metabolic pathways.

### 4.8 Importance of properties and pathways in predictive model

In order to calculate the direction of association of a property with disease, we limit each tree in the random forest to split on 3 variables. We then backtrace; if a higher value of the property led to an IBD prediction, we add one to the association score between IBD and that variable. Likewise, if a lower value of the property led to IBD, we subtract one from the association score.

In calculating metabolic pathway importance to the predictive model, we first find all properties that are consistently associated with health or with IBD. Then, we count the number of times each pathway is significantly correlated with one of those properties. If a pathway is significantly correlated with more than two consistently predictive properties, it is considered important in that phenotype.

### 4.9 Dataset

Embeddings were trained using data from the American Gut Project (24). This crowdsourced project provides 16S samples from the United States, United Kingdom, and Australia, along with associated dietary, lifestyle, and disease diagnosis information. Amplicon Sequences Variants (ASVs) were called using the DADA2 algorithm (64), resulting in 18,750 samples and 335,457 ASVs. Samples with fewer than 5,000 reads and ASV’s not occurring in at least. 07 percent of samples (13 samples) were then discarded, resulting in 18,480 samples and 26,726 ASVs. Embeddings were trained on a randomly selected 85 percent of the filtered samples, and the other 15 percent were set aside for classifier testing.

Training embeddings does not require labeled data, and so samples could be used irrespective of their available metadata. The machine learning classifier was trained and tested only on samples that had a positive or negative IBD diagnosis, 5018 and 856 samples respectively. IBD diagnosis was provided in various self-reported options from the American Gut study: “I do not have this condition”, “Self-diagnosed”, “Diagnosed by a medical professional (doctor, physician assistant)”, or “diagnosed by an alternative medicine practitioner”. For this study, we considered only samples claiming a medical professional diagnosis to be true.

Lastly, in order to test the generalizability of embeddings, we used 16S data on patients with Crohn’s Disease (CD) and Ulcerative Colitis (UC) and healthy controls from Halfvarson et. al (8). DADA2 (64) was again used to call ASVs, samples were discarded if they had fewer than 10,000 reads, and ASVs were not filtered for prevalence. After quality control, 26,251 ASVs remain, 17,775 of which have near neighbor representations in embedding space. The dataset included samples with multiple diagnoses, but for the sake of consistency, we focused on the most common diagnoses of Crohn’s disease, Ulcerative Colitis, and healthy control. In total, this left 564 samples from 118 patients, as the dataset contains multiple timepoints for each patient. When models were trained and tested on Halfvarson datasets, timepoints from the same patients were included entirely in the train or test set, so as not to train then test on the samples from the same patient.

### 4.10 Machine Learning Performance Metrics

We used two main performance metrics: area under the Receiver Operating Curve (AUROC) and area under the Precision-Recall Curve (AUPR). The Receiver Operating Curve plots true positive calls against false positive calls. The higher the AUROC, the more confident you can be that a positive prediction by the classifier is correct. The Precision-Recall Curve plots the precision, how confident you are that a positive call is correct, against recall, how many of the positive samples in the dataset were identified. A high AUPR means the classifier is able to identify most of the positive samples without making too many false positive calls. For both metrics, a value of 1 is a perfect classifier.

### 4.11 Workflow

The workflow is as described in Figure 1: First, we learn the embedding space using taxa-taxa co-occurrence data from the American Gut Project (A). The data contains 18,480 samples and 26,726 ASVs. Two taxa are considered cooccurring if they are detected in the same fecal sample. From the patterns of co-occurrence across all samples, the GloVe algorithm produces a transformation matrix, where each Amplicon Sequence Variant (taxa) is represented by a vector in embedding space (B). We call each dimension in embedding space a “property” (*P*_1_…*P*_*k*_) as each is a set of numbers used to differentiate taxas’ co-occurrence patterns. No metadata is used to create the embeddings; the process is completely unsupervised. To transform the dataset of interest into embedding space (E), we take the dot product between the dataset (D) and the transformation matrix (C). The dot product operation outputs a matrix of samples by properties, where property vectors are calculated as the average of property vectors over all the taxa present in that sample.

Lastly, we input the transformed data into a random forest classifier (F), along with 13 sample-associated features like exercise frequency, probiotic consumption, frequency of vegetable intake (G), to train a model that predicts IBD vs. Healthy host status. Samples and their associated features can be found in Suppl. Table 5.

In total, three random forest classifiers are trained, with the three types of input data: GloVe embedded, PCA transformed, and non-embedded normalized count data. Each classifier was cross-trained on 85 percent of samples to optimize hyperparameter choices for the number of decision trees, the depth of each tree, and the weight put on a positive classification.

### 4.12 Software and packages

Python Packages: Pandas 0.23.4, Numpy 1.16.3, Sklearn 0.20.2, Scipy 1.2.0, Matplotlib 3.0.0, Re 2.2.1, Skbio 0.5.5

R packages: pheatmap 1.0.12, cowplot 0.9.4, ggplot2 3.2.0, RColorBrewer 1.1-2, Gtools 3.8.1, Dada2 1.10.1, Rcpp 1.0.1, plyr 1.8.4, stylo 0.6.9, KEGGREST 1.22.0

## 5 Acknowledgements

Thanks to the Piphillan Development Team at Second Genome, for running the appropriate version of Piphillan to align ASVs to the KEGG Database, to Nathan Waugh for manuscript editing, and to the Center for Genomics and Computational Research at Oregon State University for providing computational resources and technical support.

## Notes

#### Summary of Updates

Abstract updated to clarify, Figures place in line with text, Acknowledgements updated

https://github.com/MaudeDavidLab/embeddings_final

